# tealeaves: an R package for modelling leaf temperature using energy budgets

**DOI:** 10.1101/529487

**Authors:** Christopher. D. Muir

**Affiliations:** Department of Botany, University of Hawai’i, Honolulu, Hawai’i 96822, USA

**Keywords:** boundary layer, energy balance, leaf size, leaf temperature, mathematical model, plant leaves, plant physiology, R

## Abstract

Plants must regulate leaf temperature to optimize photosynthesis, control water loss, and prevent damage caused by overheating or freezing. Physical models of leaf energy budgets calculate the energy fluxes and leaf temperatures for a given set leaf and environmental parameters. These models can provide deep insight into the variation in leaf form and function, but there are few computational tools available to use these models. Here I introduce a new R package called **tealeaves** to make complex leaf energy budget models accessible to a broader array of plant scientists. This package enables novice users to start modelling leaf energy budgets quickly while allowing experts customize their parameter settings. The code is open source, freely available, and readily integrates with other R tools for scientific computing. This paper describes the current functionality of **tealeaves**, but new features will be added in future releases. This software tool will advance new research on leaf thermal physiology to advance our understanding of basic and applied plant science.

**Summary:** **tealeaves** is a new R package to implement complex, customizable leaf energy budget models as part of an open source, transparent workflow.

## Introduction

Organisms closely regulate temperature because temperature influences many biological processes. Plants grow, survive, and reproduce under a wide variety of temperatures because natural selection endows them with adaptations to cope with different thermal regimes. Cushion plants in the alpine grown near the ground to warm up, desert plants decrease absorptance to cool down (Ehleringer *et al*., 1976), and plants keep stomata open, which can protect against extreme heat waves (Drake *et al*., 2018). These diverse mechanisms of thermal adaptation and acclimation are fascinating. Understanding them may provide insight into how plants respond to increasing temperatures and how these responses influence ecosystem function with anthropogenic climate change (Rogers *et al*., 2017). Because leaves are the primary photosynthetic organ in most plants, regulating leaf temperature is critical (Berry & Björkman, 1980). Photosynthesis peaks at intermediate temperatures(Sage & Kubien, 2007). When leaves are too warm, evaporation increases exponentially, photo- and non-photorespiratory losses subtract from carbon gain (Jones, 2014), and critical loss of function occurs about ~ 50° C (O’Sullivan *et al*., 2017). When leaves are too cold, maximum photosynthetic rates decline and can lead to damage from excess solar radiation (Huner *et al*., 1993) as well as nighttime dew and frost formation (Jordan & Smith, 1994). Natural selection should favor leaf morphologies and physiological responses that optimize leaf temperature in a given environment (Parkhurst & Loucks, 1972; Okajima *et al*., 2012; Michaletz *et al*., 2016).

To understand leaf thermal physiology, plant scientists need mathematical and computational tools to model leaf temperature as a function of leaf traits and the environment. Balancing energy budgets is a powerful mathematical tool for understanding how leaf traits and environmental parameters influnce plant physiology that has been used for over a century (Raschke, 1960). The equilibrium leaf temperature is that in which the energy gained from incoming solar and infrared radiation is balanced by that lost through infrared re-radiation, sensible heat loss/gain, and latent heat loss through transpiration (Gutschick, 2016). Leaf angle, size, and conductance to water vapour alter leaf temperature by changing how much solar radiation they intercept and how much heat they lose through sensible and latent heat loss. Likewise, enrvironmental factors such as sunlight, air temperature, humidity, and wind speed influence heat transfer between leaves and the surrounding microclimate (Gutschick, 2016). Hence, leaf energy budget models can offer deep insight on plant thermal physiology by asking how temperature is affected by one factor in isolation or in combination with another.

Leaf energy budget models have many applications, but perhaps their most widespread use is in modelling optimal leaf size and shape. The boundary layer of still air just above and below the leaf surface determines sensible and latent heat transfer and is proportional to leaf size (Gates, 1968). All else being equal, larger leaves have a thicker boundary layer, slowing heat transfer and decoupling leaf temperature from air temperature. This likely explains why, for example, many warm desert species have small leaves (Gibson, 1998). Using leaf energy budgets, Parkhurst & Loucks (1972) further predicted that leaves should be small in cold air and large under warm, shaded conditions. More recently, Okajima *et al*. (2012) extended these models, showing that small leaves maximize photosynthetic rate under high insolation and warm temperatures, but large leaves increase water-use efficiency in shadier habitats. Wright *et al*. (2017) used energy budget models to show that dew and frost formation may select against large leaves at high latitudes. Energy budget models also help explain variation in leaf shape, such as lobing and dissection, because heat transfer is determined by effective leaf width (aka characteristic leaf dimension (Taylor, 1975)) rather than total area. Effective leaf width is “the diameter of the largest circle that can be inscribed within the margin” (Leigh *et al*., 2017). Lower effective leaf width reduces leaf temperature under natural conditions in the sun (Leigh *et al*., 2017) and is under selection in sunny, drier habitats (Ferris *et al*., 2015). Besides leaf size and shape, energy balance models are useful in understanding many plant processes and traits (Gates, 1965), such as transpiration (Gates, 1968), stomatal arrangments (Foster & Smith, 1986), leaf thickness (Leigh *et al*., 2012), response to sunflecks (Schymanski *et al*., 2013), carbon economics (Michaletz *et al*., 2016), and water-use efficiency (Schymanski & Or, 2016).

Despite the utility of leaf energy budget models, there are a dearth of open source, customizable, computational tools to implement them. The **plantecophys** package implements a similar energy budget model (Duursma, 2015). However, the model is simplified for faster computation needed in ecosystem and global land surface models (Leuning, 1995). Therefore, it does not incorporate features such as different boundary layer conductances on each leaf surface, nor can users easily change default parameters for specialized cases. The Landflux website also has an Excel spreadsheet for leaf energy budgets (Tu & Fisher, 2019), but it is prohibitively time-consuming and not reproducible to use spreadsheets for large-scale simulations. Because computational tools are limited, potential users must develop models anew and learn the numerical methods necessary to find solutions. Ideally, there should be a platform in which novices can model leaf temperature to solve an interesting problem without having to write their own model and learn complicated numerical algorithms. At the same time, we need a platform that can be easily modified for experts that want to extend existing leaf energy balance models.

The goal of this study is therefore to develop software that models leaf temperature as a function of leaf traits and the environment with physical realism. This software should be open source so that the methods are transparent and code can be modified by other researchers. Secondly, it should be readily available to novice modelers yet customizable by those working on more specific problems. Finally, it should easily integrate with other advanced tools for scientific computing. To that end, I developed an R package called **tealeaves** to model leaf temepature in response to a wide variety of leaf and environmental parameters. The source code is open source and available to modify; it is easy to use with default parameters, but also customisable; and because it is written in R, the output from **tealeaves** can be analyzed and visualsized with the vast array of computational tools availble in the R environment.

## Methods

Annotated source code to generate this manuscript is available on GitHub (https://github.com/cdmuir/tealeaves-ms).

Leaf energy budgets consist of incoming radiation from solar (aka shortwave) and thermal infrared (aka longwave) sources. Leaves lose energy through infrared re-radiation, sensible heat loss, and latent heat loss due to evaporation. When leaves reach a thermal equilibrium with their environment – generally within a few minutes – these incoming and outgoing energy sources balance one another. Formally, one solves for the leaf temperature at which:

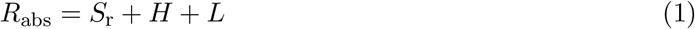

where *R*_abs_ is the absorbed radiation, *S*_r_ is infrared re-radiation, *H* is sensible heat loss, and *L* is latent heat loss. (Tables 1 and 2) lists all mathematical symbols in parameter inputs and calculated output values. Table S1 lists current default parameter values and realistic ranges with references to the literatire. This section describes the theoretical background, implementation in R, and worked examples.

**Table 1:** Parameter inputs for **tealeaves**. Each parameter has a mathematical symbol used in the text, the R character string used in the **tealeaves** package, a brief description, and the units. For physical constants, a value is provided where applicable, though users can modify these if desired.

| Symbol | R character | Description | Units |
| --- | --- | --- | --- |
| **Leaf parameters:** |  |  |  |
| $d$ | `leafsize` | leaf characteristic dimension | m |
| $\alpha_l$ | `abs_l` | absorptivity of longwave radiation (4 - 80 $\mu\text{m}$) | none |
| $\alpha_s$ | `abs_s` | absorptivity of shortwave radiation (0.3 - 4 $\mu\text{m}$) | none |
| $g_{sw}$ | `g_sw` | stomatal conductance to water vapour | $\mu\text{mol m}^{-2} \text{s}^{-1} \text{Pa}^{-1} \dagger$ |
| $g_{uw}$ | `g_uw` | cuticular conductance to water vapour | $\mu\text{mol m}^{-2} \text{s}^{-1} \text{Pa}^{-1} \dagger$ |
| $sr$ | `logit_sr` | stomatal ratio (logit transformed) | none |
| **Environmental parameters:** |  |  |  |
| $P$ | `P` | atmospheric pressure | kPa |
| $r$ | `r` | reflectance for short-wave irradiance (albedo) | none |
| $RH$ | `RH` | relative humidity | none |
| $S_{sw}$ | `S_sw` | incident short-wave (solar) radiation flux density | $\text{W m}^{-2}$ |
| $T_{air}$ | `T_air` | air temperature | K |
| $u$ | `wind` | windspeed | $\text{m s}^{-1}$ |
| **Physical constants:** |  |  |  |
| $a, b, c, d$ | `a, b, c, d` | coefficients for calculating $Nu$ and $Sh$ numbers | none |
| $c_p$ | `c_p` | heat capacity of air | $1.01 \text{ J g}^{-1} \text{K}^{-1}$ |
| $D_{h,0}$ | `D_h0` | diffusion coefficient for heat in air at 0 °C | $19.0 \times 10^{-6} \text{ m}^2 \text{s}^{-1}$ |
| $D_{m,0}$ | `D_m0` | diffusion coefficient for momentum in air at 0 °C | $13.3 \times 10^{-6} \text{ m}^2 \text{s}^{-1}$ |
| $D_{w,0}$ | `D_w0` | diffusion coefficient for water vapour in air at 0 °C | $21.2 \times 10^{-6} \text{ m}^2 \text{s}^{-1}$ |
| $\epsilon$ | `epsilon` | ratio of water to air molar masses | 0.622 |
| $eT$ | `eT` | exponent for temperature dependence of diffusion | 1.75 |
| $G$ | `G` | gravitational acceleration | $9.8 \text{ m s}^{-2}$ |
| $R$ | `R` | ideal gas constant | $8.3144598 \text{ J mol}^{-1} \text{K}^{-1}$ |
| $R_{air}$ | `R_air` | specific gas constant for dry air | $287.058 \text{ J kg}^{-1} \text{K}^{-1}$ |
| $\sigma$ | `s` | Stephan-Boltzmann constant | $5.67 \times 10^{-8} \text{ W m}^{-2} \text{K}^{-4}$ |

† conductances are presented in molar units for consistency with literature on photosynthesis but are converted to $\text{m s}^{-1}$ using the ideal gas law (see text for details) to match conductance to heat transfer.

**Table 2:** Calculated parameter and outputs for **tealeaves**. Some parameters are intermediate calculations (see Methods) but are not included in the **tealeaves** output (see R documentation accompanying package for further detail). Each parameter has a mathematical symbol used in the text, the R character string used in the **tealeaves** package, a brief description, and the units.

| Symbol | R character | Description | Units |
| --- | --- | --- | --- |
| **Leaf parameters:** |  |  |  |
| $E$ | E | transpiration rate | $\text{mol m}^{-2} \text{s}^{-1}$ |
| $g_h$ | g_h | boundary layer conductance to heat | $\text{m s}^{-1}$ |
| $g_{bw}$ | g_bw | boundary layer conductance to water vapour | $\text{m s}^{-1}$ |
| $g_{tw}$ | g_tw | total conductance to water vapour | $\text{m s}^{-1}$ |
| $Gr$ | Gr | Grashof number | none |
| $H$ | H | sensible heat loss | $\text{W m}^{-2}$ |
| $L$ | L | latent heat loss | $\text{W m}^{-2}$ |
| $Nu$ | Nu | Nusselt number | none |
| $R_{abs}$ | R_abs | absorbed radiation | $\text{W m}^{-2}$ |
| $Re$ | Re | Reynolds number | none |
| $S_r$ | S_r | infrared re-radiation | $\text{W m}^{-2}$ |
| $Sh$ | Sh | Sherwood number | none |
| $T_{leaf}$ | T_leaf | leaf temperature | K |
| **Environmental parameters:** |  |  |  |
| $d_{wv}$ | d_wv | water vapour pressure differential | $\text{mol m}^{-3}$ |
| $h_{vap}$ | h_vap | latent heat of vapourization | $\text{J mol}^{-1}$ |
| $P_a$ | P_a | density of dry air | $\text{g m}^{-3}$ |
| $p_{air}$ | p_air | water vapour pressure of the air | kPa |
| $p_{sat}$ | p_sat | saturating water vapour pressure | kPa |
| $S_{lw}$ | S_lw | incident long-wave (thermal infrared) radiation flux density | $\text{W m}^{-2}$ |
| $T_{sky}$ | T_sky | clear sky temperature | K |
| **Physical constants:** |  |  |  |
| $D_h$ | D_h | diffusion coefficient for heat in air at a given temperature | $\text{m}^2 \text{s}^{-1}$ |
| $D_m$ | D_m | diffusion coefficient for momentum in air at a given temperature | $\text{m}^2 \text{s}^{-1}$ |
| $D_w$ | D_w | diffusion coefficient for water vapour in air at a given temperature | $\text{m}^2 \text{s}^{-1}$ |
| **Convergence diagnostics:** |  |  |  |
| | value | energy balance at equilibrium $T_{leaf}$ | $\text{W m}^{-2}$ |
|  | convergence | 0 = converged; 1 = failed | none |

## Theory

This section describes the current **tealeaves** implementation. However, future releases will alter some assumptions and incorporate new features. I mention future modifications in the Discussion.

### Absorbed radiation

The **tealeaves** model for absorbed radiation follows Okajima *et al*. (2012):

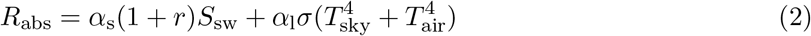

The left half of the equation calculates absorbed solar radiation; the right half includes thermal infrared radiation. As in Okajima *et al*. (2012), I calculated *T*_sky_ as a function of *T*_air_:

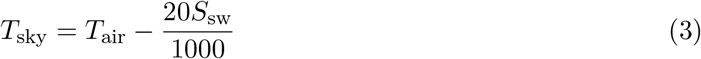

### Thermal infrared re-radiation

Both leaf surfaces reradiate thermal infrared radiation as a function of leaf emissivity (equal to the infrared absorption, *α*_l_) and air temperature (Foster & Smith, 1986; Okajima *et al*., 2012):

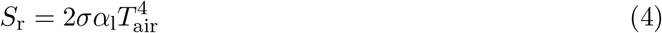

### Sensible heat flux

Sensible heat flux (*H*) is calculated as:

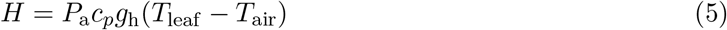

The density of dry air (*P*_a_) is calculated as in Foster & Smith (1986):

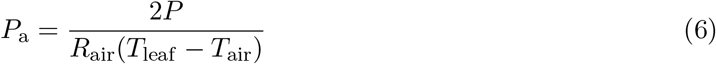

**tealeaves** sums the boundary layer conductance to heat for both the upper and lower surface following Foster & Smith (1986), assuming a horizontal leaf orientation:

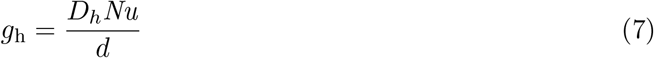

The diffusion coefficient of heat in air is a function of temperature and pressure:

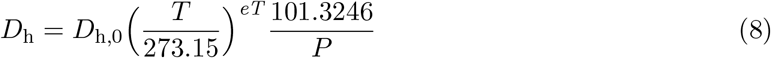

The temperature dependence of diffusion (*eT*) is generally between 1.5-2 for heat and water vapour (Monteith & Unsworth, 2013). To calculate diffusion coefficients, **tealeaves** uses the average of the leaf and air temperature: *T* = (*T*_air_ + *T*_leaf_)/2. The Nusselt number *Nu* is modeled as a mixed convection:

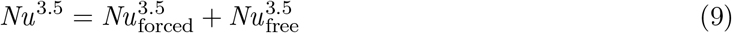

where

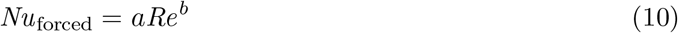

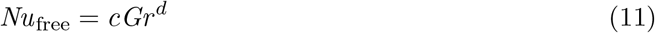

*a, b, c, d* are constants that depend on whether flow is laminar or turbulent and the direction of flow in the case of free convection (see below). In general, when the Archimedes number *Ar* = *Gr*/*Re*^2^ ≪ 0.1, free convection dominates; when *Ar* = *Gr*/*Re*^2^ ≫ 10, forced convection dominates (Nobel, 2009). The Nusselt number coefficients can be found in Monteith & Unsworth (2013). For forced convection, flow is laminar if *Re* < 4000, *a* = 0.6, *b* = 0.5; flow is turbulent if *Re* > 4000, *a* = 0.032, *b* = 0.8. These cutoffs for leaves are lower than for artificial surfaces because trichomes and other anatomical features of leaf surfaces induce turbulence more readily (Grace & Wilson, 1976). For free convection, flow is laminar. For the upper surface when *T*_leaf_ > *T*_air_ or the lower surface when *T*_leaf_ < *T*_air_, *c* = 0.5, *d* = 0.25. Conversely, for the lower surface when *T*_leaf_ > *T*_air_ or the upper surface when *T*_leaf_ < *T*_air_, *c* = 0.23, *d* = 0.25.

Grashof and Reynolds numbers are calculated as follows:

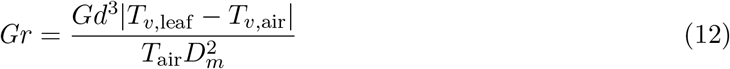

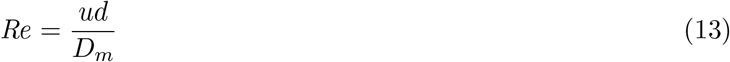

The diffusion coefficient for momentum in air (*D_m_*) is calculated for a given temperature following the same procedure above for heat diffusion (*D_h_*; see Eq. 8). The virtual temperature is calculated according to Monteith & Unsworth (2013) assuming that the leaf airspace is fully saturated while the air is has a vapour pressure decifit of *p*_air_:

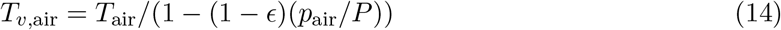

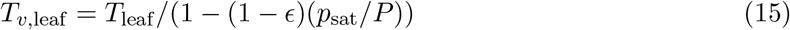

The saturation water vapour pressure *p*_sat_ as a function of temperature is calculated using the Goff-Gratch equation (Vömdel, 2016). The vapour pressure of air is calculated from the relative humidity as *p*_air_ = *RHp*_sat_.

### Latent heat flux and evaporation

Latent heat loss is the product of the latent heat of vaporization, the total leaf conductance to water vapour, and the water vapour gradient:

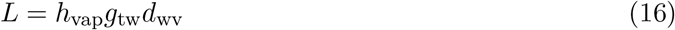

The latent heat of vapourization (*h*_vap_) is a linear function of temperature. **tealeaves** calculates *h*_vap_ using parameters estimated from linear regression on data from Nobel (2009):

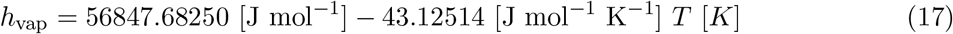

The water vapour pressure differential from the inside to the outside of the leaf is the water vapor pressure inside the leaf, which is assumed to be saturated (*p*_leaf_ = *p*_sat_), minus the water vapor pressure of the air (*p*_air_), calculated as described above. This value is converted from kPa to mol m^−3^ using the ideal gas law:

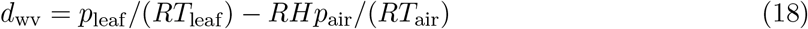

The total conductance to water vapor (*g*_tw_) is the sum of the parallel lower (usually abaxial) and upper (usually adaxial) conductances

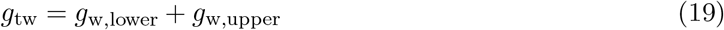

The conductance to water vapor on each surface is a function of parallel stomatal (*g*_sw_) and cuticular (*g*_uw_) conductances in series with the boundary layer conductance (*g*_bw_). The stomatal, cuticular, and boundary layer conductance on the lower surface are:

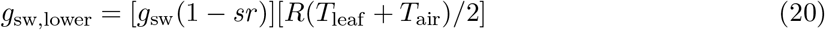

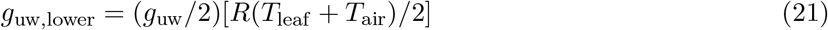

Note that the user provides the total leaf stomatal and cuticular conductance to water vapur in units of *μ*mol m^−2^ s^−1^ Pa^−1^, which are then converted to units of m s^−1^ using the ideal gas law. Stomatal conductance is partitioned among leaf surfaces depending on stomatal ratio (*sr*); cuticular conductance is assumed equal on each leaf surface. The corresponding expressions for the upper surface are:

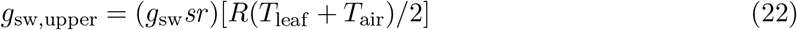

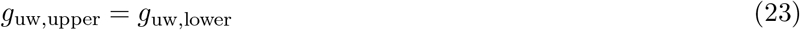

The boundary layer conductances for each surface differ because of free convection (Foster & Smith, 1986) and are calcualted similarly to that for heat (Eq. 7):

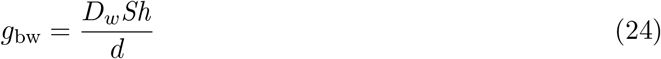

*D_w_* is calculated using the Eq. 8, except that is *D*_*w*,0_ is substituted for *D*_*h*,0_. Each surface has its own Sherwood number (*Sh*):

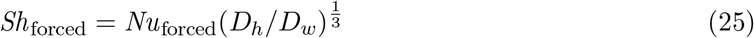

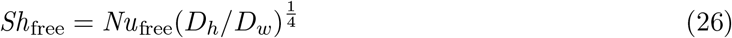

As with *Nu*, *Sh* is calculated assuming mixed convection:

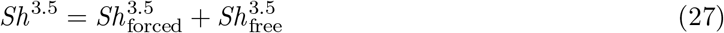

Evaporation rate (mol H_2_O m^−2^ s^−1^) is the product of the total conductance to water vapour (Eqn 19) and the water vapour gradient (Eqn 18):

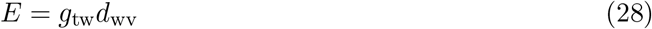

### Solving in R

R is a fully open source programming language for statistical computing that allows users to develop their own packages with new functions. **tealeaves** takes three sets of parameter inputs: leaf parameters, environmental parameters, and physical constants (see Table 1). The package provides reasonable defaults, but users can input new values to address their question, as I demonstrate in the next section. With one or more parameter sets, **tealeaves** uses the `uniroot` function in R base package **stats** to find the *T*_leaf_ that balances the leaf energy budget (Eqn 1). It outputs the equilibrium *T*_leaf_ and energy fluxes in a table for analysis and visualization.

Unlike previous leaf energy models, **tealeaves** ensures that calculations are technically correct by assigning stadard SI units with the R package **units** Pebesma *et al*. (2016). Every parameter and calculated value must have correctly assigned units. If units are not properly defined, **tealeaves** will produce an error because it is unable to convert values. To ensure accuracy, these unitless functions are tested against their counterparts with units using the **testthat** package (Wickham, 2011). Other R packages that contributed to **tealeaves** are **crayon** (Csárdi, 2017), **dplyr** (Wickham *et al*., 2018), **glue** (Hester, 2018), **furrr** (Vaughan & Dancho, 2018), **future** (Bengtsson, 2018), **ggplot** (Wickham, 2016), **magrittr** (Bache & Wickham, 2014), **purrr** (Henry & Wickham, 2018a), **rlang** (Henry & Wickham, 2018b), **stringr** (Wickham, 2018), **tidyr** (Wickham & Henry, 2018).

### Worked examples

In this section, I provide two worked examples; more complex worked examples are found in the Supporting Information. The first illustrates that it is straightforward to use **tealeaves** with a few lines of code with default settings. The second shows that it is also possible to model *T*_leaf_ across multiple leaf trait and environmental gradients for more advanced applications.

**Example 1: a minimum worked example**

The box below provides R code implementing the minimum worked example with default settings.

~~~
>  library(tealeaves)
>  # Default parameter inputs
>  leaf_par   <- make_leafpar()
>  enviro_par <- make_enviropar()
>  constants  <- make_constants()
>  # Solve for T_leaf
>  T_leaf <- tleaf(leaf_par, enviro_par, constants,
+                  quiet = TRUE)
>
~~~

**Example 2: leaf temperature along environmental gradients**

The box below provides R code to calculate leaf temperature along an air temperature gradient for leaves of different sizes.

~~~
>  library(tealeaves)
>  # Custom parameter inputs
>  leaf_par   <- make_leafpar(
+    replace = list(
+      leafsize = set_units(c(0.0025, 0.025, 0.25), “m”)
+      )
+  )
>  enviro_par <- make_enviropar(
+    replace = list(
+      T_air = set_units(seq(275, 310, 5), “K”)
+      )
+  )
>  constants <- make_constants()
>  # Solve for T_leaf over a range of T_air
>  T_leaves <- tleaves(leaf_par, enviro_par, constants,
+                      quiet = TRUE)
>
~~~

### Extended examples

To see the range of possible applications for **tealeaves**, I ran four additional sets of simulations. The first models the leaf-to-air temperature differential for different leaves sizes across a gradient of air tempuratures; the second models the leaf-to-air temperature differential across a gradient of incident solar radiation for different stomatal conductances; the third models the leaf-to-air temperature differential for different sized leaves under free, mixed, and forced convection; and the fourth models the effect of stomatal ratio on evaporation under free and forced convection. These extended examples are documented more fully in the Supporting Information with accompanying R code.

To provide a sense of which leaf and environmental parameters affect *T*_leaf_ the most under “typical” conditions, I varied *g*_sw_, *d*, sr, *RH, S*_sw_, and *u* over a wide range of realistic values while holding all other values constant at their default setting (Table S1).

## Results

### tealeaves’s source code is open to all

A development version of **tealeaves** is currently available on GitHub (https://github.com/cdmuir/tealeaves). A stable version of **tealeaves** will be released on the Comprehensive R Archive Network (CRAN, https://cran.r-project.org/) after peer-review to ensure that the underlying model has been vetted by expert plant scientists. I will continue developing the package and depositing revised source code on GitHub between stable release versions. Other plant scientists can contribute code to improve **tealeaves** or modify the source code on their own installations for a more fully customized implementation.

### tealeaves is straightfoward to use and modify

**tealeaves** lowers the activation energy to start using leaf energy budgets in a transparent and reproducible workflow. Default settings provide a reasonable starting point (see Worked Example 1 and Table S1), but they should be carefully inspected to ensure that are appropriate for particular questions. At default settings, low stomatal conductance, high humidity, and/or low wind speed cause leaf temperatures to heat substantially above air temperature (Fig. S1A,D,F). Small leaves are closely coupled to air temperature, whereas large leaves are not (Fig. S1B). Leaves can operate below air temperature at low light, but above it at higher light (Fig. S1E). Stomatal ratio has only a modest effect on leaf temperature (Fig. S1C). Most users will want to modify these settings and simulate leaf temperature over a range of leaf and environmental parameters, so these results are not generalizable to all cases. By design, **tealeaves** easily allows users to define multiple simultaneous trait and environmental gradients (see Worked Example 2 and Fig. 1).

**Figure 1:**
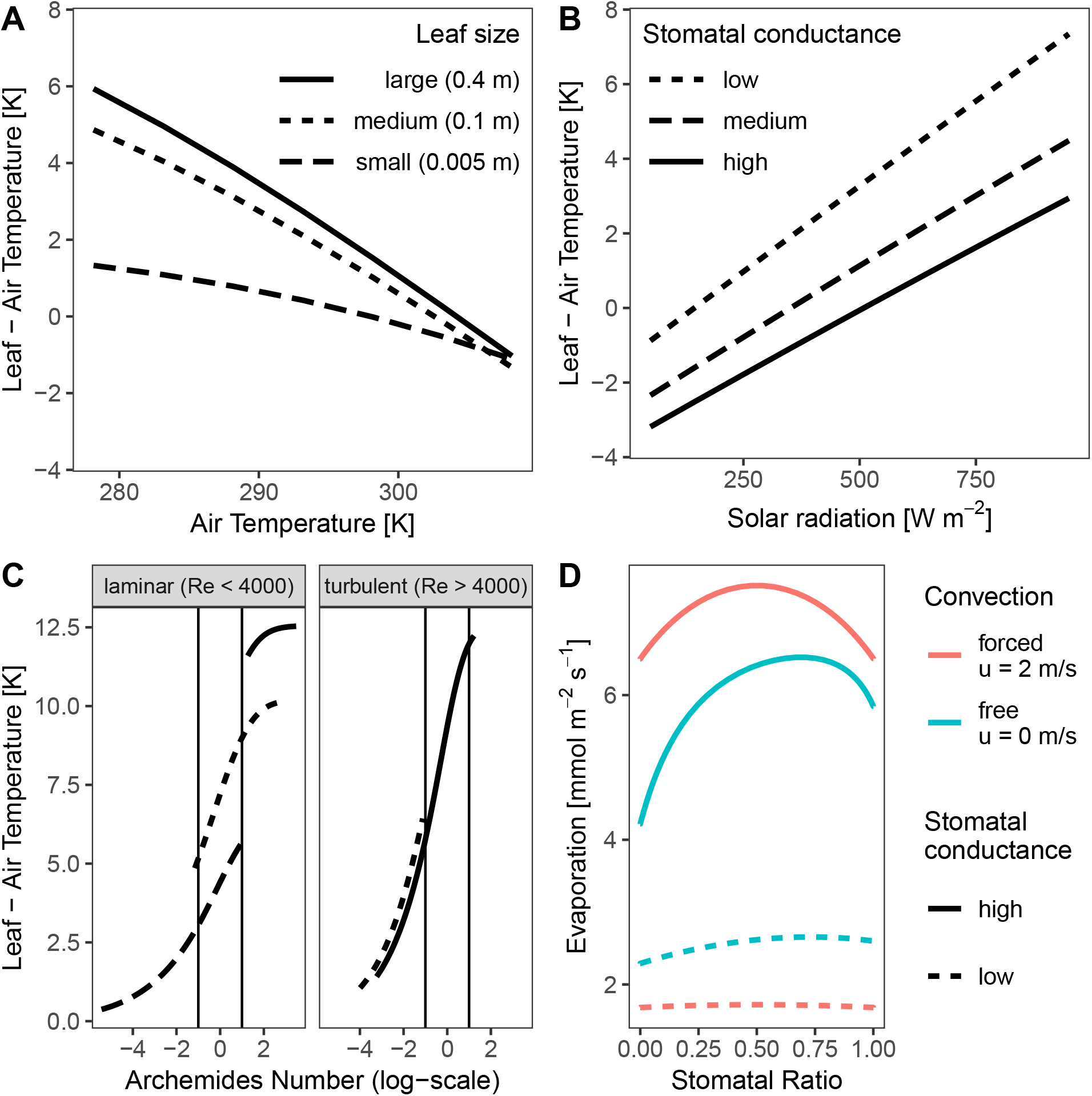
Extended examples of **tealeaves**. Code to generate these examples is provided in the Supporting Information. A. The temperature of smaller leaves is more closely coupled to air temperature. Each line represents a different leaf size (small, dashed line; medium, dotted line; large, solid line) and the leaf-to-air temperature differential (*y*-axis) over an air temperature gradient (*x*-axis). B. Greater stomatal conductance cools leaves. Each line represents a different stomatal conductance (low, dashed line, 1 *μ*mol m^−2^ s^−1^ Pa^−1^; medium, dotted line, 3 *μ*mol m^−2^ s^−1^ Pa^−1^; high, solid line, 5 *μ*mol m^−2^ s^−1^ Pa^−1^) and the leaf-to-air temperature differential (*y*-axis) over a gradient of incident solar radiation (*x*-axis). C. Forced convection dominates in small leaves; free convection dominates in very large leaves. Leaf size is indicated by line type as in Panel A. Vertical lines indicate approximate shifts from forced convection (*Ar* < 0.1), mixed convection (0.1 < *Ar* < 10), and free convection (*Ar* > 10). Small leaves always experience forced convection, leading to lower leaf temperature compared to large leaves experiencing free convection. D. Amphistomatous leaves (Stomatal Ratio ~ 0.5) evaporate more than hypo- or hyperstomatous leaves (Stomatal Ratio ~ 0 or 1, respectively), especially under free convection (low wind speed, *u*).

## Discussion

Scientists have used energy budgets to model leaf temperature for over a century (see Raschke [1960] for historical references). Despite many advances in our understanding of the environmental and leaf parameters that affect heat exchange (Gutschick, 2016), there exist few computational tools to implement complex energy budget models. The **tealeaves** package fills this gap by providing platform for modelling energy budgets in a transparent and reproducible way with R (R Core Team, 2018), a freely available and widely used programming language for scientific computing. Unlike previous software, **tealeaves** removes ambiguity by forcing users to specify proper SI units through the R package **units** (Pebesma *et al*., 2016). Neophytes with little experience modelling leaf temperature may get started quickly without having to develop their model *de novo*, while specialists can modify the open source code to customize **tealeaves** to their specificiations. **tealeaves** also readily integrates with the vast array of data analysis and visualization tools in R. These features will enable wider adoption of leaf energy budgets models to understand plant biology. However, as I discuss below, the current version of **tealeaves** has several important limitiations that can be addressed in future releases.

**tealeaves** provides a computational platform for beginners and experts alike to model leaf temperature using energy budgets. Previously, researchers wanting to implement sophisticated leaf energy budget models that required numerical solutions had to write their model and learn a numerical algorithm to solve it. Most often, these solutions are not published and/or are not open source. This slows down research for nonspecialists by introducing unnecessary barriers and can be error-prone. For example, the current **tealeaves** model relies on previous work by Foster & Smith (1986). Without a platform like **tealeaves**, extending their work required developing the mathematical and computational tools *de novo* every time. Also, the published version Foster & Smith (1986) contains several small errors and tyographical inconsistencies in the equations. While these are most likely mistakes made during typesetting and publication, without open source code, it is very challenging to determine if these mistakes also occurred in their computer simulations. Transparent, open source code does not prevent mistakes, but makes it easier for the community to discover mistakes and fix them faster.

Ultimately, the goal of **tealeaves** is to provide a platform for implementing realistic and fully customizable energy budget models. Such models may take too much computational time to be useful for large-scale ecosystem models, but they can help understand a wider range of fascinating and poorly understood leaf anatomical and morphological features, as well as identify under what conditions simpler leaf temperature models are adequate. The Introduction lists several possible uses, but most of these problems cannot currently be solved with **tealeaves** alone. For example, many photosynthetic processes are temperature sensitive, but it would require simultaneous modelling of leaf temperature, stomatal conductance, and photosynthesis to predict optimal trait values. **tealeaves** should therefore be thought of as one component in an expanding ecosystem of interrelated tools for modelling plant physiology. A standalone package that only models leaf temperature is best suited for flexible integration with existing and yet-to-be-developed tools.

Currently, **tealeaves** has several limitations that I plan to address in future releases. It uses rather simple models of infrared radiation and direct versus diffuse radiation. Ideally, it would be better if users could supply their own functions to calculate these parameters from the total irradiance. The model also assumes leaves are horizontal, whereas leaf orientation varies widely. Following previous authors, I modeled heat transfer as a mixed convection (Eqns 9 and 27, but this may not adequately describe real leaf heat exchange (Roth-Nebelsick, 2001). **tealeaves** calculates equilibrium as opposed to transient behavior, which may takes several minutes to reach. Finally, the model assumes a single homogenous leaf temperature rather than using finite element modelling to calculate leaf temperature gradients across leaves of different shapes. These are important limitations of the current software which can be addressed in future work.

In conclusion, **tealeaves** provides an open source software platform for leaf energy balance models in R. Leaf energy balance models are highly useful tools for understanding plant form and function and new computational tools will make these models more broadly accessible, advancing basic and applied plant science.

## Acknowledgements

Tom Buckley kindly explained how to convert conductance from molar to ‘engineering’ units.

## Supporting Information

### Supporting Tables

**Table S1:** Reasonable values for **tealeaves** parameter inputs with references to the primary literature. The current version of **tealeaves** uses a default value within the range of reasonable values. See Table 1 for a key to symbols.

| Symbol | **tealeaves** Default | Range | Reference(s) |
| --- | --- | --- | --- |
| **Leaf parameters:** |  |  |  |
| $d$ | 0.1 | 0.004 – 0.4 m | Wright *et al.* (2017) |
| $\alpha_l$ | 0.97 | 0.95 – 0.97 | Gutschick (2016) |
| $\alpha_s$ | 0.5 | 0.4 – 0.6 | Jones (2014) |
| $g_{sw}$ | 5 | 0.01 – 10 $\mu\text{mol m}^{-2} \text{s}^{-1} \text{Pa}^{-1}$ | Lin *et al.* (2015); Duursma *et al.* (2019) |
| $g_{uw}$ | 0.1 | 0.01 – 1 $\mu\text{mol m}^{-2} \text{s}^{-1} \text{Pa}^{-1}$ | Duursma *et al.* (2019) |
| $SR$ | 0.5 | 0 – 1 (untransformed) | Muir (2015) |
| **Environmental parameters:** |  |  |  |
| $P$ | 101.3246 | 50 (5000 mas) – 100 (0 mas) kPa | Körner (2007); Stull (2011) |
| $r$ | 0.2 | 0.1 (lava) – 0.6 (ice) | Stull (2011) |
| $RH$ | 0.5 | 0 – 1 | Jones (2014) |
| $S_{sw}$ | 1000 | 0 – 1000 $\text{W m}^{-2}$ | Jones (2014) |
| $T_{air}$ | 298.15 | 270 – 320 K | Jones (2014) |
| $u$ | 2 | 0 – 10 $\text{m s}^{-1}$ | Vogel (2009) |

### Supporting Figures

**Figure S1:**
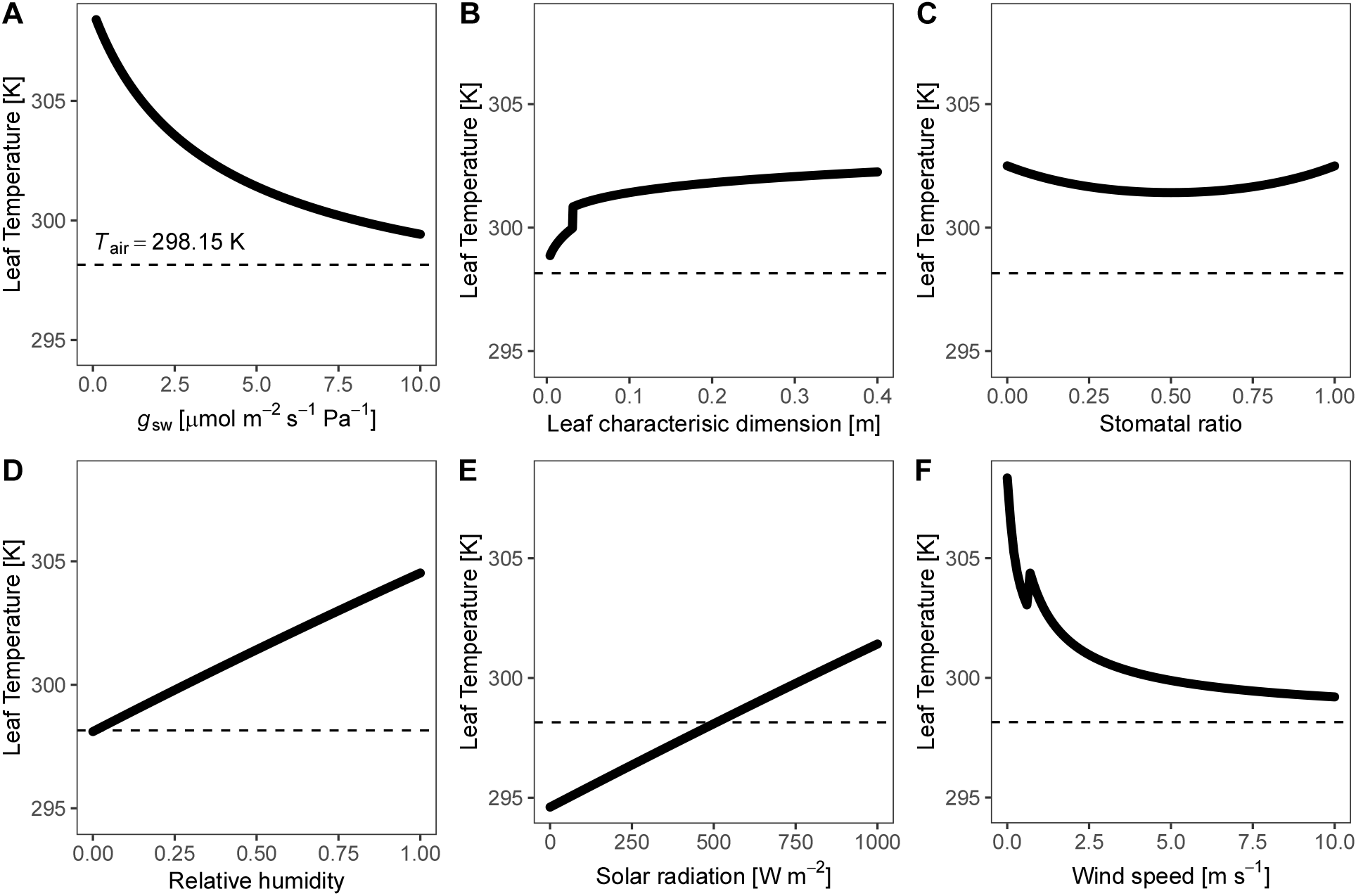
The effect of key leaf (**A – C**) and environmental (**D – F**) parameters on leaf temperature, holding other parameters constant. **A**) Greater stomatal conductance (*g*_sw_, *x*-axis) reduces leaf temperature through latent heat loss. **B**) Larger leaves (*d, x*-axis) have thicker boundary layers, causing them to heat up more in the sun. **C**) Amphistomatous leaves (*SR* = 0.5, *x*-axis) lose more water through transpiration than leaves with all stomata on one surface, leading to a lower leaf temperature. **D**) Greater humidity (*RH, x*-axis) increases leaf temperature by limiting latent heat loss. **E**) With low solar radiation (*S*_sw_, *x*-axis), leaf temperature is below air temperature; with high solar radiation, leaf temperature is greater than air temperature. **F**) At greater wind speeds (*u, x*-axis) leaf temperature is more closely coupled to air temperature. The discontinuity represents the shift from laminar to turbulent flow. For reference, the dashed line is the air temperature in all simulations. All calculations used the following leaf parameter values unless they varied: *d* = 0.1 m; *α*_s_ = 0.5; *α*_l_ = 0.97; *g*_sw_ = 5 *μ*mol m^−2^ s^−1^ Pa^−1^; *g*_uw_ = 0.1 *μ*mol m^−2^ s^−1^ Pa^−1^; *SR* = 0.5. All calculations used the following environmental parameter values unless they varied: *P* = 101.3246 kPa; *r* = 0.2; *RH* = 0.5; *S*_sw_ = 1000 W m^−2^; *T*_air_ = 298.15 K; *u* = 2 m s^−1^. See Table 1 for symbol definitions.

### Extended examples

R code for running extended examples (Fig. 1). The below code and the code to generate figures are deposited on GitHub (https://github.com/cdmuir/tealeaves-ms).

~~~
> # Extended example 1:
> # leaf size and leaf-to-air temperature differential
>
> library(tealeaves)
> lp <- make_leafpar(
+   replace = list(
+    leafsize = set_units(c(0.005, 0.1, 0.4), “m”)
+   )
+ )
> ep <- make_enviropar(
+   replace = list(
+    S_sw = set_units(660, “W/m^2”),
+    T_air = set_units(seq(278.15, 308.15, 5), “K”)
+   )
+ )
> exe1 <- tleaves(lp, ep, cs, progress = TRUE, quiet = TRUE,
+                 set_units = TRUE, parallel = TRUE)
~~~

~~~
> # Extended example 2:
> # Solar radiation and leaf-to-air temperature differential
>
> library(tealeaves)
> lp <- make_leafpar(
+   replace = list(
+     g_sw = set_units(c(1, 3, 5), “umol/m^2/s/Pa”)
+   )
+ )
> ep <- make_enviropar(
+   replace = list(
+     S_sw = set_units(seq(50, 950, 100), “W/m^2”)
+  )
+ )
> exe2 <- tleaves(lp, ep, cs, progress = TRUE, quiet = TRUE,
+                 set_units = TRUE, parallel = TRUE)
>
~~~

~~~
> # Extended example 3:
> # Wind speed and leaf-to-air temperature differential
>
> library(tealeaves)
> lp <- make_leafpar(
+   replace = list(
+     leafsize = set_units(c(0.005, 0.1, 0.5), “m”)
+   )
+ )
> ep <- make_enviropar(
+   replace = list(
+     wind = set_units(exp(seq(log(0.01), log(10),
+                     length.out = 1e2)), “m/s”)
+   )
+ )
> exe3 <- tleaves(lp, ep, cs, progress = TRUE, quiet = TRUE,
+                 set_units = TRUE, parallel = TRUE)
~~~

~~~
> # Extended example 4:
> # Stomatal ratio and evaporation
>
> library(tealeaves)
> lp <- make_leafpar(
+   replace = list(
+     g_sw = set_units(c(0.4, 4), “umol/s/m^2/Pa”),
+     logit_sr = set_units(seq(−10, 10, length.out = 1e2))
+   )
+ )
> ep <- make_enviropar(
+   replace = list(
+     RH = set_units(0.2),
+     T_air = set_units(293.15, “K”),
+     wind = set_units(c(0, 2), “m/s”)
+   )
+ )
> exe4 <- tleaves(lp, ep, cs, progress = TRUE, quiet = TRUE,
+                 set_units = TRUE, parallel = TRUE)
>
~~~

